# Expression and function of murine WFDC2 in the respiratory tract

**DOI:** 10.1101/2020.05.05.079293

**Authors:** L Bingle, H Armes, DJ Williams, O Gianfrancesco, Md M K Chowdhury, R Drapkin, C D Bingle

## Abstract

*WFDC2/HE4* encodes a poorly characterised secretory protein that shares structural similarity with multifunctional host defence proteins through possession of two conserved Whey Acidic Protein/four disulphide-core (WFDC) domains. WFDC2 is expressed in multiple epithelia and although its’ function remains unresolved, it is also overexpressed in a number of human cancers and has an established role as a cancer marker. Currently, little is known about the distribution of WFDC2 in the mouse and thus we have systematically analysed the mouse *wfdc2* gene, its’ expression and distribution. We have used recombinant WFDC2 for functional studies. *Wfdc2* is the most highly expressed family member in the lung and is enriched in the nasopharynx. *Wfdc2* is the most highly expressed family member in differentiated epithelial cells isolated from the trachea, nasal passages and middle ear. *Wfdc2* consists of 5 exons with exon 3 encoding an unstructured linker region that separates the two WFDC domains. This genomic organisation appears to be restricted to the *Muridae* and *Cricetidae* families of rodents. Similar to the situation in man, mouse *wfdc2* can be alternatively spliced to yield a number of distinct transcripts that have the potential to generate a repertoire of distinct protein isoforms. We used immunohistochemistry to localise the proteins to tissues of the respiratory tract and head and neck regions. Although the protein was limited to epithelial cells of the respiratory tract and nasal and oral cavities, it was expressed in different cells in different regions suggesting expression is governed by a unique regulatory mechanism. Recombinant WFDC2 did not possess antiproteinase activity against trypsin or elastase and had no clear antimicrobial activity.

## Introduction

The whey acidic protein/four disulphide core (WFDC) domain is a conserved protein fold containing 8 highly conserved cysteine residues which form a characteristic 4 disulphide bond motif [1]. Although the domain is found throughout metazoans most research has focused on the proteins in mammals, reptiles, crustaceans and molluscs. Most WFDC domains are found in small secretory molecules that display a variety of functions [2,3]. In some cases they exist as the sole structural domain in a protein, whereas other proteins may contain multiple domains perhaps mixed with other defined folds and some are found in proteins that are much larger [3]. Small secreted WFDC containing proteins make up a small family that exhibit hallmarks of rapid molecular evolution [3–5]. The majority of the genes encoding these proteins are localised in a single genomic loci, interspersed with a number of other genes. On chromosome 20 in man the locus contains 14 genes [4–7] whereas in mice a locus on chromosome 2 contains 16 genes [8]. The two best-studied members of this family are the antiproteinase/alarmin proteins, Secretory Leukocyte Protease Inhibitor (SLPI)/WFDC4 and elafin/PI3/WFDC14 [9, 10]. These proteins display a pleiotropic range of functions and are important host defence molecules. Both are secreted from epithelial surfaces and have been shown to be important in host defence in the lung. PI3 is not found in mice [8]. The other WFDC proteins have generally been suggested to function in a similar way, largely in a “guilt by association” manner [3,9,10] but there remains limited functional information about many of these genes and often information is restricted to expression data. The relative importance of WFDC domain containing proteins in the respiratory tract is unknown [11].

We have previously shown that the double WFDC domain containing protein, WFDC2 is highly expressed in the respiratory tract and shown that its expression is increased in the lungs of patients with severe cystic fibrosis lung disease [12,13]. WFDC2 was initially identified as a transcript highly expressed in the epididymis [14], hence is was originally called Human epididymis 4 (HE4), and it has been widely studied as a secreted cancer marker, initially in ovarian cancer [15,16] but subsequently in many other tumours [17–19]. Some evidence suggests that WFDC2 may be a marker of other diseases [20,21] and functions in innate defence of the gut [22]. Recently, two studies have unexpectedly shown that loss of WFDC2 leads to a respiratory insufficiency and embryonic lethality [23,24]. The driver of this phenotype remains unresolved.

In the present study we investigated the expression of *WFDC* genes in respiratory tissues of the mouse. On the basis that WFDC2 was by far the most highly expressed gene in this region we have performed a systematic analysis of *Wfdc2*, analysed its’ expression and alternative splicing, studied its distribution by immunohistochemistry (IHC) and undertaken preliminary functional analysis.

## MATERIALS AND METHODS

### Sample collection

C57BL6 mice of mixed sex were used in this study. All animal experiments were performed in accordance with the UK Animals (Scientific procedures) Act, authorized under a UK Home Office License, and approved by the animal project review committee of the University of Sheffield. We collected tissues for RNA extraction and IHC as described [25]. Samples from the nasal passages were isolated as described [26]. Some tissues for IHC were collected following perfusion fixation and all samples were fixed in cold 4% paraformaldehyde. Where required, decalcification of mouse heads was performed using formic acid. Fixed tissues were embedded in paraffin according to standard protocols.

### mTEC, mNEC and mMEEC cell cultures

Mouse tracheal, nasal and middle ear epithelial cell cultures were established from C57BL/6J mice and cultured at the air-liquid interface to generate well-differentiated cultures (mTEC, mNEC and mMEEC) as previously described [27, 28]. Five or six mice were used for each cell isolation. mTEC and mNEC cells were grown at the ALI for 14 days and mMEEC cells were cultured for 7 days. Differentiated ALI cultures displayed phenotypic features associated with a complex population of cells that matched with the features of the native epithelium [27,28]. At the end of the culture period, RNA was isolated using the Direct-zol system (Zymo Research). Apical washes of secretory proteins were collected by washing the surface of the differentiated cultures with 100ul of PBS.

### Clariom S mouse microarray

Samples of total RNA were checked for quantity and quality using the Nanodrop and Agilent Bionalyser 600. Subsequently 200ng was prepared for analysis on the Clariom S mouse GeneChip (Thermo Fisher) according to the manufacturers’ instructions. Briefly mRNA was converted to double stranded cDNA with the incorporation of a T7 polymerase binding site at the 3’ end of the RNA molecule. Antisense RNA was generated by utilization of the T7 polymerase. This was purified using a magnetic bead process and further quantified on the Nanodrop. 15ug of the aRNA was taken forward to generate a sense DNA strand. This was fragmented and end labelled with biotin before being incorporated into a hybridization solution and incubated with the GeneChip. Following post hybridisation washing using the fluidics station a fluorescent signal corresponding to hybridization of the labelled material to the oligonucleotide probes on the chip was achieved using a streptavidin-phycoerythrein cocktail. Gene chips were scanned on the Gene Chip 7000G scanner and the images collected as CEL files. Microarray data in CEL files was analysed using Affymetrix Expression Console. Gene level summary data were obtained using the RMA summarization method. Date was Log2 transformed for generation of relative gene expression levels.

### Bioinformatic analysis and molecular characterisation of mWFDC2

The RefSeq coding sequence of mouse WFDC2 (NP_080599.1) was used to search the NCBI non-redundant protein database to identify representative protein sequences from across mammalian phylogeny. Obvious potential discrepancies in extracted peptide sequences were adjusted by subjecting the genomic sequence surrounding the WFDC2 gene location to gene prediction analysis by GENESCAN (http://hollywood.mit.edu/GENSCAN.html). Multiple sequence alignment was performed using Clustal Omega (http://www.ebi.ac.uk/Tools/msa/clustalo/) and the resulting alignment was visualised using the BoxShade server (http://www.ch.embnet.org/software/BOX_form.html). Alternative splicing of mouse ESTs was identified using the BLAT server on the UCSC Genome Bioinformatics server (https://genome.ucsc.edu).

Total RNA was isolated from mouse tissues using TRI-reagent. Reverse transcription (RT) reactions were performed using oligo dT primed RNA reverse transcribed from total RNA. Initial RT-PCR was performed with primers designed to amplify within the predicted coding region of mWFDC2 (RT Forward: 5’ AGG TGG ACA GCC AGT GTT CT 3’; RT Reverse 5’ GAC CAG GAA GAA ATG CAA 3’). Subsequently, we used primers located in the 5’ and 3’ non-coding exons to identify alternative splicing that utilised these two exons, Exon 1 Forward (5’-CACCATGCCTGCCTGTCG-3’) and Exon 5 Reverse (5’-CAGAGAGAAAGGAGGCCACA-5’). PCR products were directly cloned into TOPO pCRII (Invitrogen) and sequenced for verification.

### Immunohistochemistry

Immunostaining was performed on 4µm paraffin sections mounted onto glass slides (SuperFrost Plus, VWR international, Belgium) following established protocols with antigen retrieval being performed using tri-sodium citrate buffer. Optimal antibody dilutions were determined empirically and the final dilution used was 1:500. Duplicate sections were always used as no primary antibody controls. A Vectastain Elite ABC kit (Vector Laboratories) containing a goat anti-rabbit biotin-labelled secondary antibody was used according to the manufacturer’s instructions. Peroxidase enzymatic development was performed using a Vector NovaRed substrate kit resulting in red staining in positive cells. Sections were counterstained with haematoxylin, dehydrated to xylene and mounted in DPX.

### Generation and characterisation of mouse specific WFDC2 antibodies

Antibodies against murine Wfdc2 were generated using three distinct polypeptides covering different regions of the protein: Ac-CPISATGTDAEKPGE-amide (amino acids 22-35); Ac-CPNGPSEGELSGTDTKLSE-amide (amino acids 73-90); Ac-KPPAVTREGLGVREKQGTC-amide (amino acids 114-132). The polypeptides were HPLC purified. Rabbits were immunized with all three peptides using standard procedures. Crude sera and protein A purified antibodies were evaluated after multiple boosts by ELISA (against peptides) and immunohistochemistry using cells and tissue from a murine model of ovarian cancer [29].

The antibody was validated on endogenous and recombinant samples. We used samples of apical washes recovered from fully differentiated mTEC and mNEC cells as well as recombinant protein produed as described below. Samples were run on 12.5% SDS-PAGE gels and western blotted using the WFDC2 antibody (1:2000 dilution) and detected as described [25,30]

### Functional analysis of WFDC2

Full-length murine *wfdc2* was amplified from lung RNA using primers incorporating an in-frame FLAG tag immediately preceding the stop codon. The fragment was cloned into pVR1255 and fully sequenced prior to use. The calcium phosphate transfection of HEK293 cells was used to generate recombinant protein. Following transfection cells were cultured in serum-free media for 48 hours and conditioned media collected for up to 6 days post transfection. WFDC2 protein was purified using batch absorbance on a FLAG affinity resin. Media was incubated with the resin for 2 hours and then FLAG-tagged WFDC2 was eluted using FLAG peptide buffer (150 ng/μl). Protein was analysed by Western blotting and quantified by BCA assay.

To establish whether WFDC2 is capable of inhibiting the serine proteases, trypsin, the effect on enzyme activity was monitored by measuring chromogenic substrate digestion via the subsequent accumulation of cleaved p-nitroanilide (p-NA). WFDC2 protein (0, 100, 200, 500 and 1000 nM) was pre-incubated with a fixed concentration of trypsin for 15 minutes to allow for binding between the potential inhibitor and enzyme substrate. Chromogenic substrate was added and the reaction was incubated for 15 minutes at 37°C. The reaction was quenched and the absorbance measured at 405 nm. Recombinant human SLPI (0, 100, 200, 500 and 1000 nM) and a broad-spectrum protease inhibitor (PI) solution (added in sequentially increasing volumes) were used as controls for the assay.

The effect of WFDC2 on bacterial viability was also monitored on *Streptococcus mutans, Streptococcus gordonii, Streptococcus pneumoniae* and *Escherichia coli* by measuring changes in absorbance of bacterial suspension over time to produce a growth curve. Briefly, bacterial culture was inoculated with WFDC2 protein (1.0 μM) and seeded into a 96 well plate. The optical density of each well was measured at 600 nm every 30 minutes for 12 hours using a microplate reader.

## RESULTS

### Expression of WFDC domain containing gene in the respiratory tract

We extracted expression data on the 16 *wfdc* genes localised chromosome 2 in lung tissue of C57/BL6 mice from LungGENS (Figure 1A). This showed that *wfdc2* was by far the most highly expressed gene in mouse lungs. *Slpi* was the next most highly expressed with the remain genes being seen at a very low level. We next studied expression of *Wfdc2* in the GeneAtlas GNF1M data set on BioGPS (Figure 1B) [31]. This showed that Wfdc2 is highly expressed not only in lung and trachea, but also in other tissue from the nasopharynx. Lower levels of expression were seen in other tissue including the ovary, uterus, bladder, kidney and GI tract. This data confirms that WFDC2 exhibits a level of tissue specificity. This tissue distribution is similar to that seen in man, where again high expression is seen in the respiratory tract [12,13]. WFDC2 expression starts to be seen in the mouse lung from e11.5 (Figure S1). We generated microarray data from cultures of mTEC, mNEC and mMEECs differentiated at the air liquid interface. This culture technique produces epithelial cultures that recapitulate the phenotype of the respective tissues [27,28]. Again, the data confirmed that *Wfdc2* is the most highly expressed *Wfdc*-family member in epithelial cells from all three tissues (Figure 1C-1E). *Slpi* was again the second most highly expressed gene.

**Figure 1:**
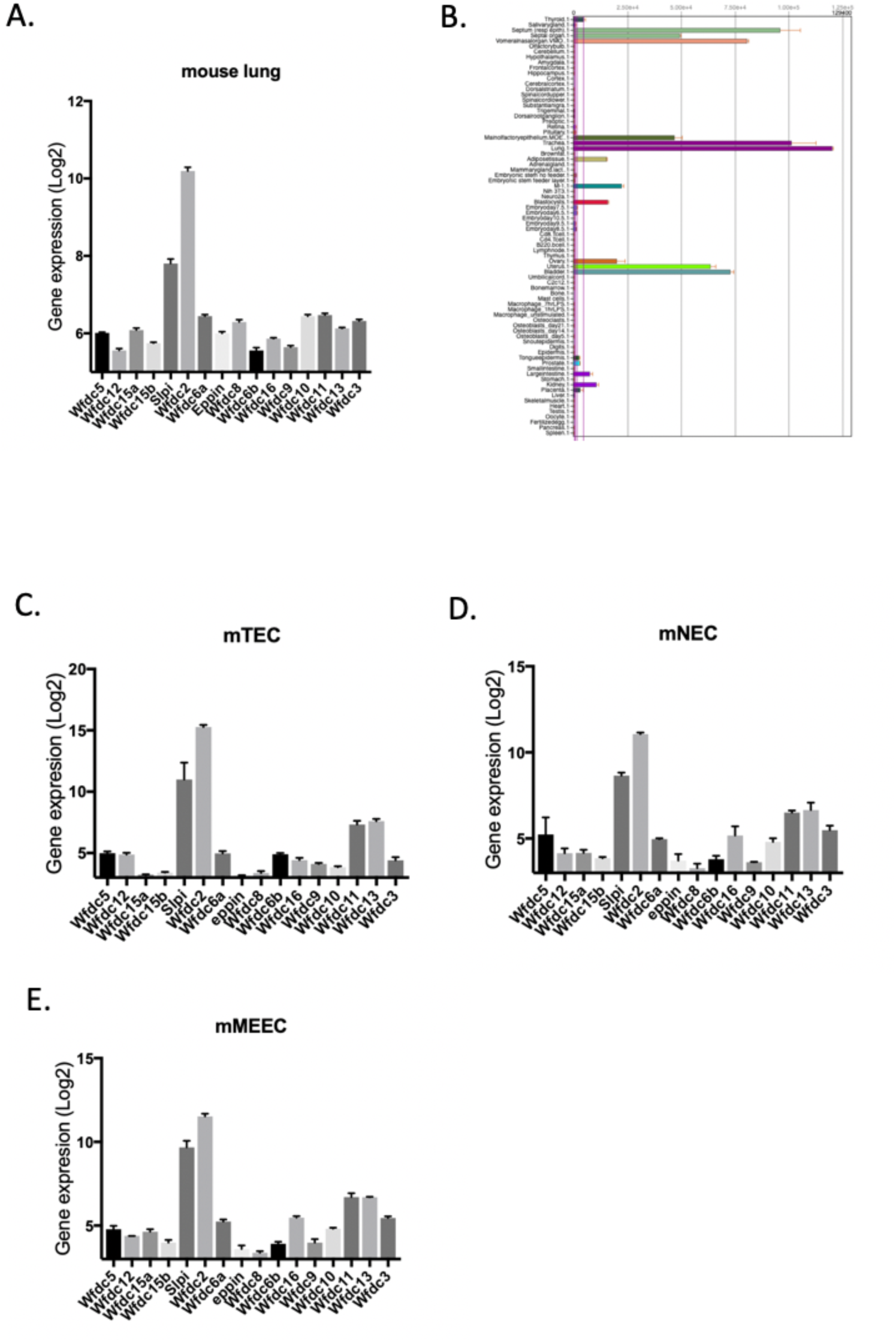
*Wfdc2* is the most highly expressed WFDC-domain containing gene in the mouse respiratory tract. **A**. Expression data for the Chromosome 2 *Wfdc* genes in C57/BL6 mouse lungs was extracted from array data in LungGENs and gene expression plotted s Log2 expression data using prism. Data is presented as mean + SEM (n=12). **B.** Expression of Wfdc2 in 78 mouse tissues/cell types from the GeneAtlas GNF1M array data set from BioGPS. **C-E.** Expression data for the Chromosome 2 *Wfdc* genes in mouse tracheal (**C**), nasal (**D**) and middle ear (**E**) epithelial cell cultures was extracted from array data generated as outlined in the Material and Methods section. Data is presented at Log2 means + SEM (n=2-3 cultures per cell type.

### Molecular characterisation and bioinformatic analysis of mouse WFDC2

We have previously shown that mouse and rat WFDC2 proteins differ from those of other species through the inclusion of an inserted region located between the two WFDC2 domains [12]. We performed BLAST analysis of the non-redundant protein database to investigate the evolutionary origins of this additional region of sequence. Protein sequences of WFDC2 from 16 separate species were aligned (Figure 2). This shows that five sequences contain a portion of sequence of between 40-53 amino acid residues in length that creates an unstructured ‘linker’ region between the two WFDC domains. Comparison of these sequences against genomic reference sequence confirmed that each was due to the use of an additional coding exon. The use of this sequence appears to be limited to the closely related *Muridae* and *Cricetidae* families of the *Eumuroida* clade of rodents, as it is not seen in the predicted protein from *Nannospalax galili*, a blind mole rat from the *Splacidae* family (Figure 2). This suggests that an insertion event likely occurred prior to the divergence of the most recent common ancestor of the two sub-families.

**Figure 2:**
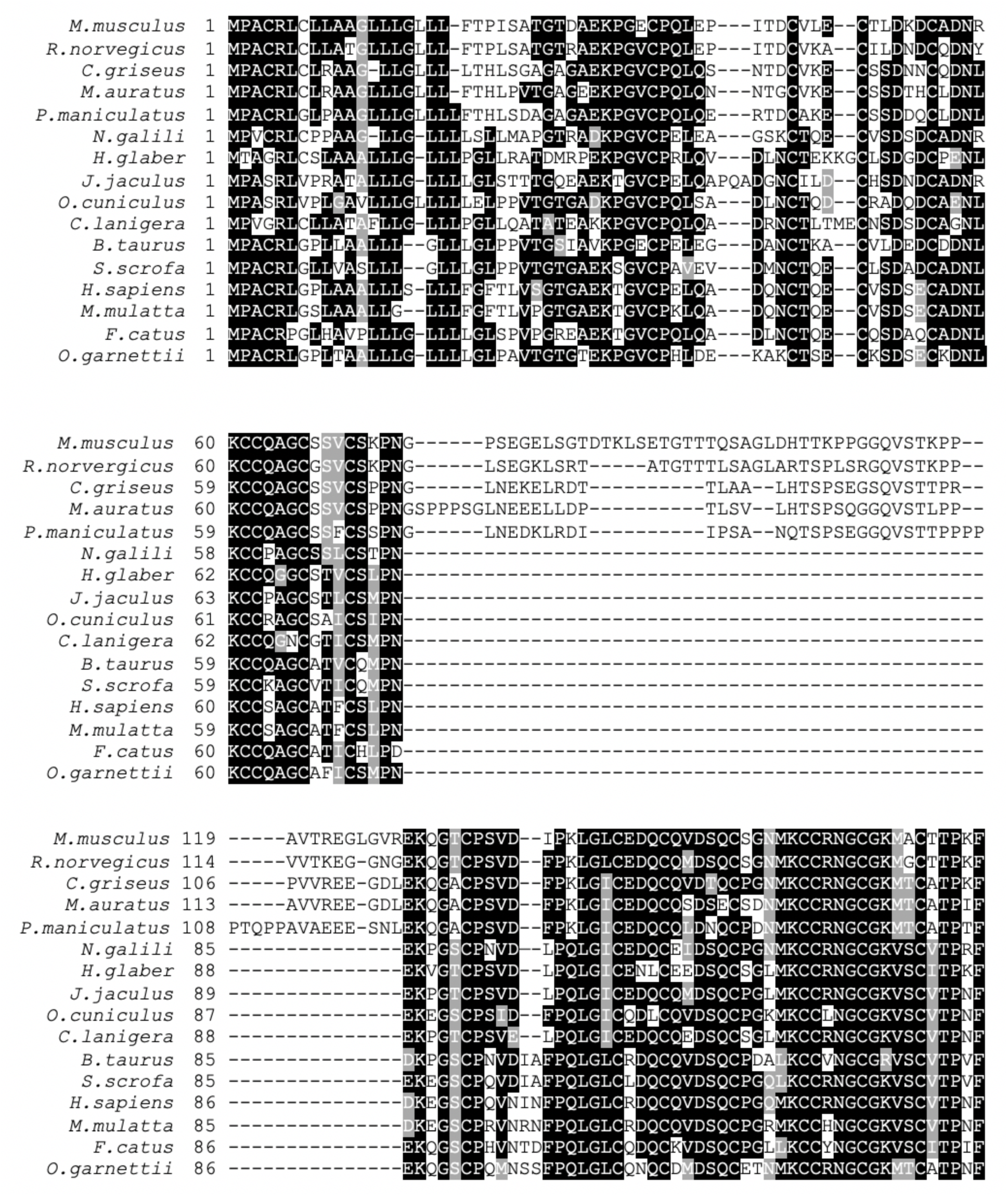
Multiple Sequence Alignment of WFDC2 sequences identifies a unique region of inserted sequence in some rodent proteins. A multiple sequence alignment of selected WFDC2 sequences was performed using Clusta Omega as described in the Materials and Methods section. The final alignment was generated using Boxshade. Species within the *Muridae* and *Cricetidae* sub-families of the *Muroidea* family contain a ~50 amino acid residue insert between the two WFDC domain regions, suggesting that it arises from a recent common ancestor,

Aside from this striking observation, the eight cysteine residues that provide the disulphide bonds that form each of the two WFDC2 domains are completely conserved between species. Overall the second WFDC domain is more highly conserved between species, being 70% identical to the same domain in the human protein, in comparison to the N terminal domains, which share 56% sequence identity. Despite the C-terminal WFDC2 domains showing this higher overall similarity, proteins from outside of the wider rodent family also include two additional amino acids in the region between cysteine residues 1 and 2 (Figure 2).

### Genomic organisation and expression of mWFDC2

The genomic organisation of *Wfdc2* is shown in figure 3A. As can be seen the gene consists of 5 exons with exons two and four encoding the N-terminal and C-terminal WFDC domains respectively. Alignment of the sequence against mouse ESTs using the UCSC BLAT server, suggested that this organisation represents the major transcript but that additional transcripts do exist.

**Figure 3.**
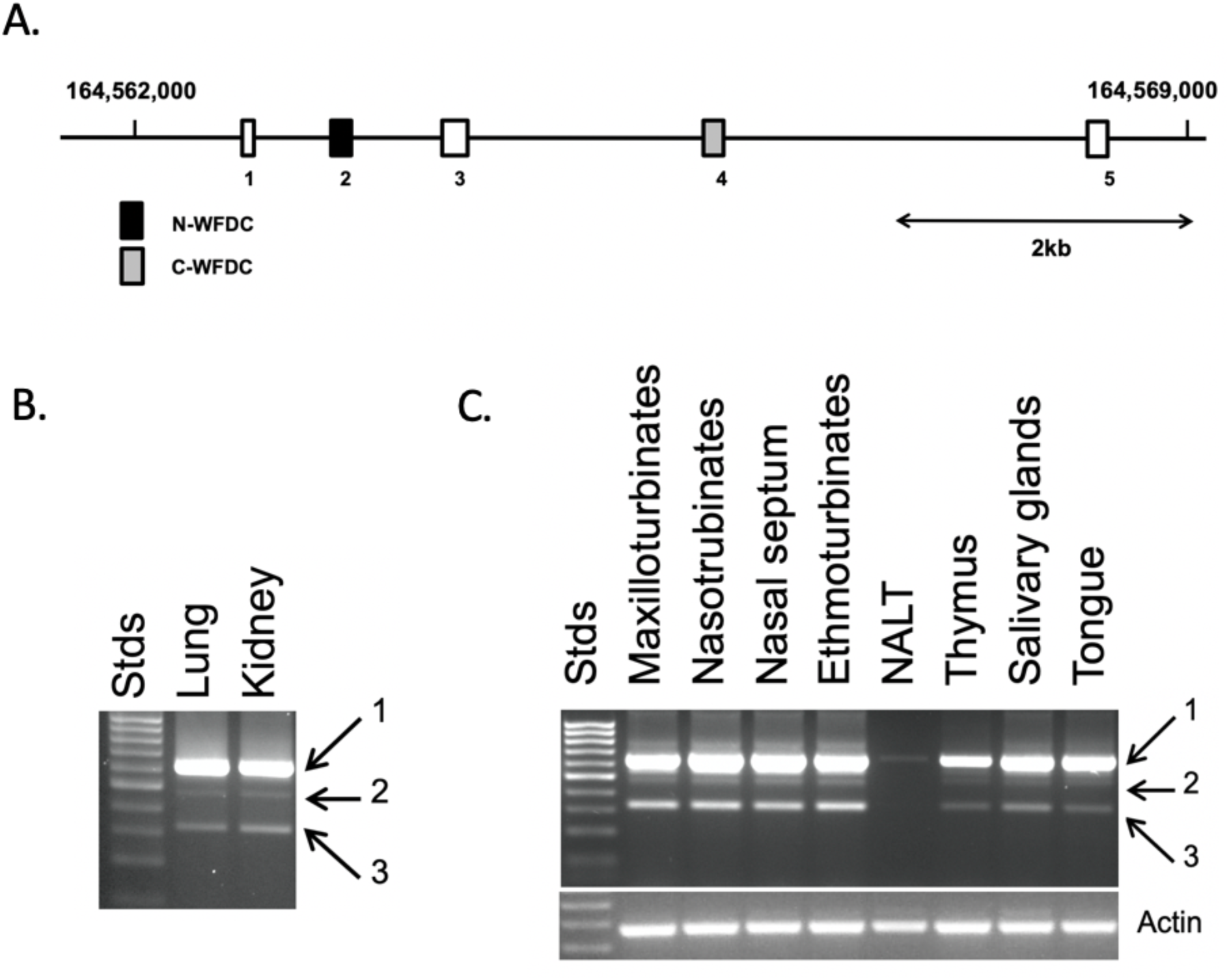
Genomic organisation and expression of mouse WFDC2 gene. **A.** Schematic representation of the *WFDC2* gene with the exon numbers shown below the figure. The WFDC domains are shaded. The numbers represent the position on the chromosome in bp. The scale bar represents 2Kb. **B.** Expression of *WFDC2* was investigated in lung and kidney RNA by RT-PCR as described in the Materials and Methods section. Primers (in exons 1 and 5) were designed to amplify the entire coding region. The position of the molecular mass markers is indicated to the side of the image. Arrows 1, 2 and 3 indicate the location of the different bands that were excised from the gel and cloned for verification of sequences. **C.** Alternative splicing of *WFDC2* was investigated in tissues from regions of the head and neck by RT-PCR as above. The position of the molecular mass markers is indicated to the side of the image. Arrows 1, 2 and 3 indicate the alternatively spliced transcripts.

Given the different gene structure seen in the mouse compared to human, and in light of our previous observation that the human WFDC2 gene undergoes alternative splicing [12], we examined alternative splicing of the gene in lung and kidney by PCR using primers located in exon 1 and exon 5. Both tissues examined generated a band of the expected size (619bp) corresponding to a transcript containing exons 1-5 (Figure 3B). This was the most abundant band in all of the samples investigated. Bands from the regions of the gel marked 2 and 3 were excised, cloned and sequenced. This confirmed that the presence of transcripts corresponding to exons 1, 2, 4, 5 (potentially yielding a protein similar to WFDC2 from other species) and exons 1, 3, 4, 5 (region 2) and exons 1, 4, 5 (region 3). To further investigate expression in tissues from the head and neck regions additional PCRs, with the same primer pair, were undertaken. Again, the band representing the wild type transcript was the most abundant in the tissues tested (Figure 3C). Strong expression of WFDC2 was seen in samples from the nasal passages (the turbinates and septum) and the tongue but there was limited expression in the nasal associated lymphoid tissue (NALT) and the extra-orbital lacrimal gland (EOLG). In all cases the most abundant of the additional bands corresponded to a transcript containing exons 1, 4 and 5. Analysis of EST sequences in the NCBI database confirmed that sequences corresponding to transcripts containing exons 1,4,5 (accession number CA494779), 1,2,4,5 (accession numbers CA482028 and AV501895) and 1, 2, 5 (BX639871) had previously been identified. The latter transcript removes the stop codon (found in exon 4) and potentially adds an additional 45 amino acids to the C-terminal of the protein isoform, ASASSSLRNGERLFLPDCASGVVPVASFLSGLCISSWSDESISFF. We did not identify a transcript corresponding to exons 1,2,3,5 in this analysis.

### Localisation of WFDC2 in mouse tissues

Having identified high gene expression for WFDC2 in the respiratory tissues and in the head and neck we produced an antibody against murine WFDC2 to study the distribution of WFDC2 in mouse tissues. The antibody was validated by western blotting with apical washes from mTEC and mNEC cells as well as recombinant protein produced in HEK cells. The antibody recognised endogenous WFCD2 in apical washes from both mTEC and mNEC cells as well as that produced in HEK cells. The antibody identified a number of immunoreactive bands ranging in size between 30-40kDa whereas the expected size of the protein is 18kDa. In the case of the recombinant WFDC2 in spite of the fact that the epitope tag adds 34 amino acids to the proteins (approximately 3.5 kDa) its molecular mass appeared to be smaller than that of the endogenous protein. It is likely that the different bands correspond to glycosylation variants and that this also accounts for the smear of protein in the apical wash from the mTEC cells.

**Figure 4.**
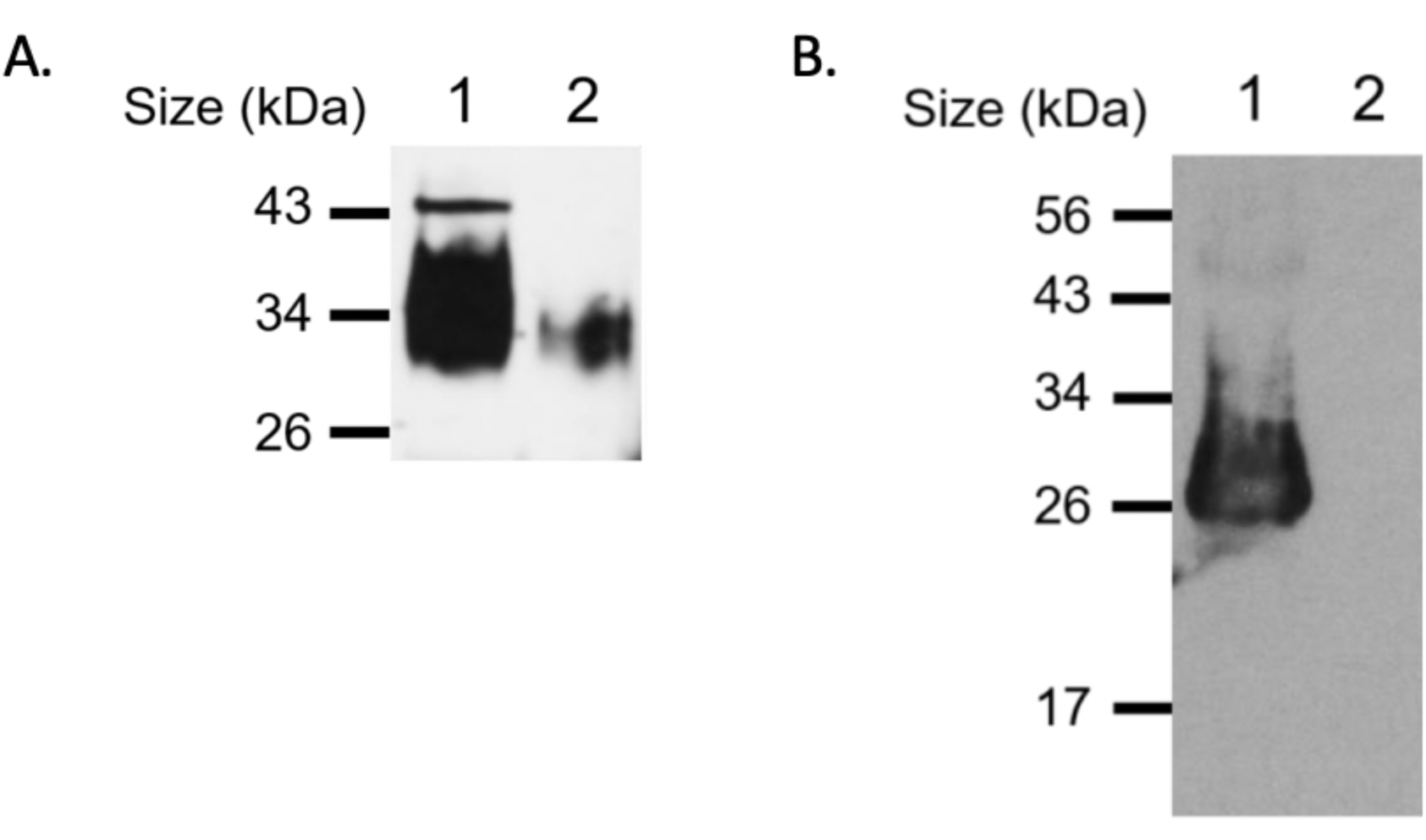
Characterisation of the WFDC2 antibody by western blotting. **A.** 20ul of apical washes from differentiated mTEC (lane 1) and mNEC (lane 2) cells, **B** and 20 of conditioned media from mWFDC2 expressing (lane 1) or mock transfected cells (lane 2) were resolved on a 12% SDS-PAGE gels. Western blots were developed using the WFDC2 polyclonal antibody (1:2000) as described in the Materials and Methods section. The black lines indicate the positions of the molecular mass markers.

Having established that the antibody could robustly detect WFDC2, we then used it to localise the protein in a range of tissues. Given that mouse *wfdc2* is most abundantly expressed in the respiratory tract and in tissues associated with the head and neck, we performed a systematic analysis of protein localisation by immunohistochemistry on tissues from these regions. In sections of lung tissue robust protein staining was observed in the epithelial cells of the bronchiolar epithelium (Figure 5A and B), with continuous staining occurring down to the level of the terminal bronchioles and to the bronchioalveolar duct junction. Peripheral lung tissue was essentially negative using this technique (Figure 5B). Moving higher up the airways staining was seen in an epithelial cell population in the trachea as well as in the serous cells of the airway submucosal glands (Figure 5C). Staining was also seen in the squamous epithelium of the epiglottis (Figure 5D) as well as in the respiratory epithelium of the eustachian tube (Figure 5E), the nasal cavity (Figure 5F), the lower portion of the nasopharynx (Figure 5G), the nasal septum (Figure 5H) and in the nasal turbinates (Figure 5I). The olfactory epithelium did not stain for the protein, but rather staining was seen in the Bowman’s (olfactory) glands underlying the olfactory epithelium and within the ducts leading to the epithelial surface (Figure 5J). Strong staining was also seen in goblet cells in the conjunctiva (Figure 5K). It was clear from these sections that WFDC2 was not restricted to a single cell type, being seen in squamous epithelium as well as in distinct epithelial cell types in the respiratory tract and nasopharynx, including goblet cells and club cells. This lack of cell-type specific localisation is also seen in scRNAseq data from whole mouse lungs (Figure S2) which shows that the gene is enriched in multiple epithelial cell types. The majority of minor glands, including the lateral nasal glands and glands of the nasal septum (Figure 5H), were also largely negative, with the exception of weak staining in the serous portion of the glands guarding the eustachian tube (Figure 5E) and within the proximal tongue (results not shown). Sections of submandibular gland exhibited strong staining in the mucus tubules (Figure 5L, M). In contrast in the parotid and sublingual glands staining was seen in the ducts and was absent from the serous and mucous acini, respectively (Figure 5L, M).

**Figure 5.**
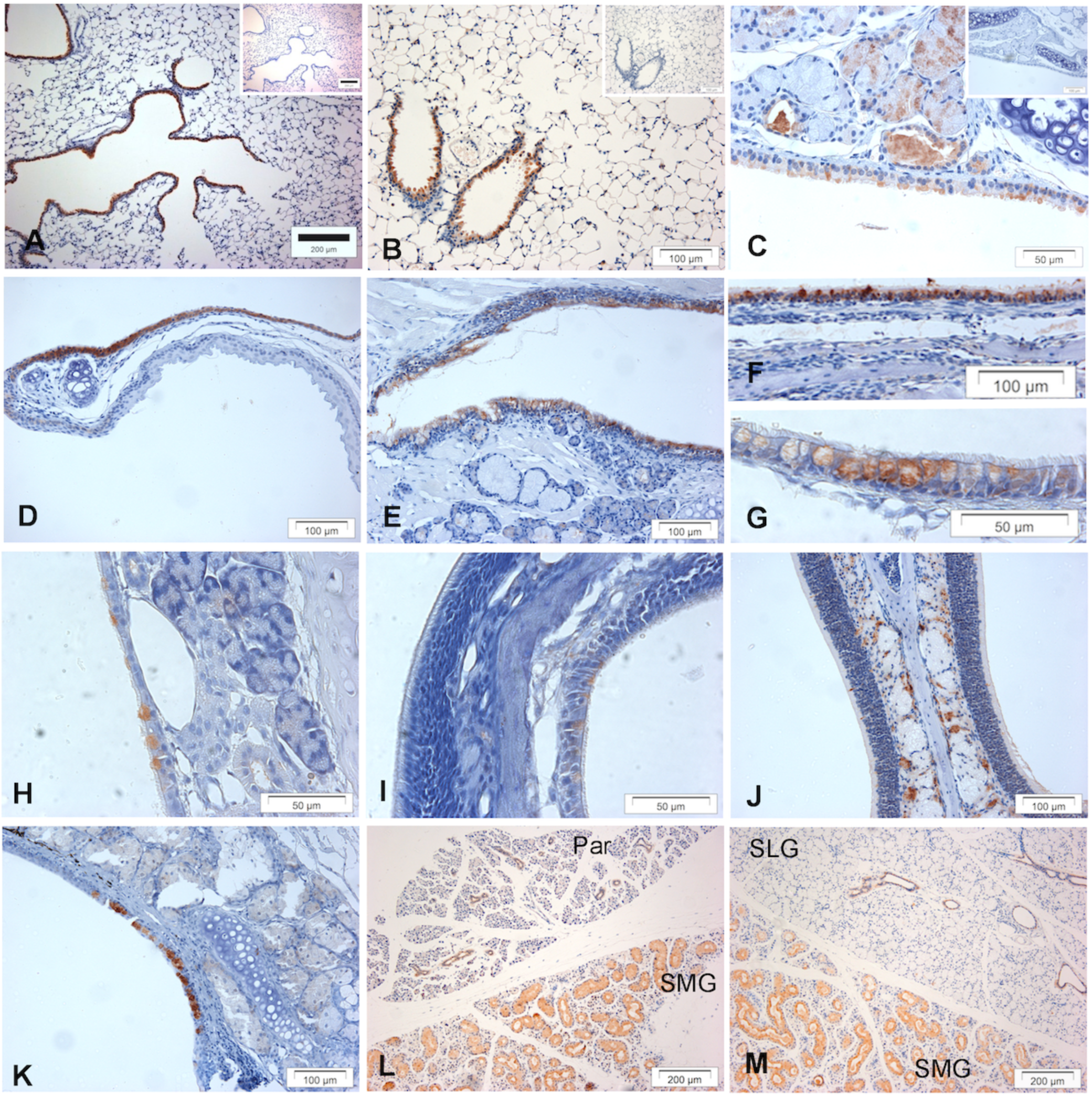
Distribution of WFDC2 in the respiratory tract, and regions of the head and neck. Immunohistochemistry for WFDC2 was performed as described in Materials and Methods section. Tissues stained were lung (**A, B**), Tracheal epithelium and submucosal gland (**C**), epiglottis (**D**) eustachian tube (**E**) the roof of the nasopharynx (**F**), tissues of the nasal turbinates and septum (**H, I, J**), the conjunctiva (**K**) and the major salivary glands (**L, M**). The individual salivary glands are indicated on the figures; Parotid gland (Par), Submandibular gland (SMG) and Sublingual Gland (SLG). The insets in panels **A**, and **C** represent negative control sections developed without primary antibody. The original magnifications of the images were ≈100X (**A, L, M**), 200X (**D, E, F, J** and **K**) and 400X (**B**, **C, G, H, I**).

### WFDC2 does not possess proteinase inhibitory or antimicrobial activities

We produced Flag-tagged mWFDC2 from transfected HEK293 cells for functional analysis. Secreted WFDC2 appeared predominantly as a smeared band from ~26 to 34 kDa. A second ~50 kDa band could be seen which potentially represents a protein dimer. Transfected cells continued to secrete protein up to 6 days post-transfection (Figure 6A). We purified WFDC2 protein from transfected HEK293 conditioned media using FLAG affinity resin. The amount of protein was significantly enriched compared to the original media sample (Figure 6B). As the recombinant protein resolved as a smeared band of 26-34 kDa, we analysed the protein for potential glycosylation sites. Unlike the human protein, mouse WFDC2 has no predicted N-glycosylation sites but it is predicted to have 17 O-glycosylation sites, 15 of which are found in exon 3. When murine WFDC2 was incubated with sialidase and analysed by Western blotting there was a visible shift in molecular weight between the digested and non-digested forms (Figure 6C), although the apparent size was still greater than would be expected from the primary amino acid sequence.

**Figure 6:**
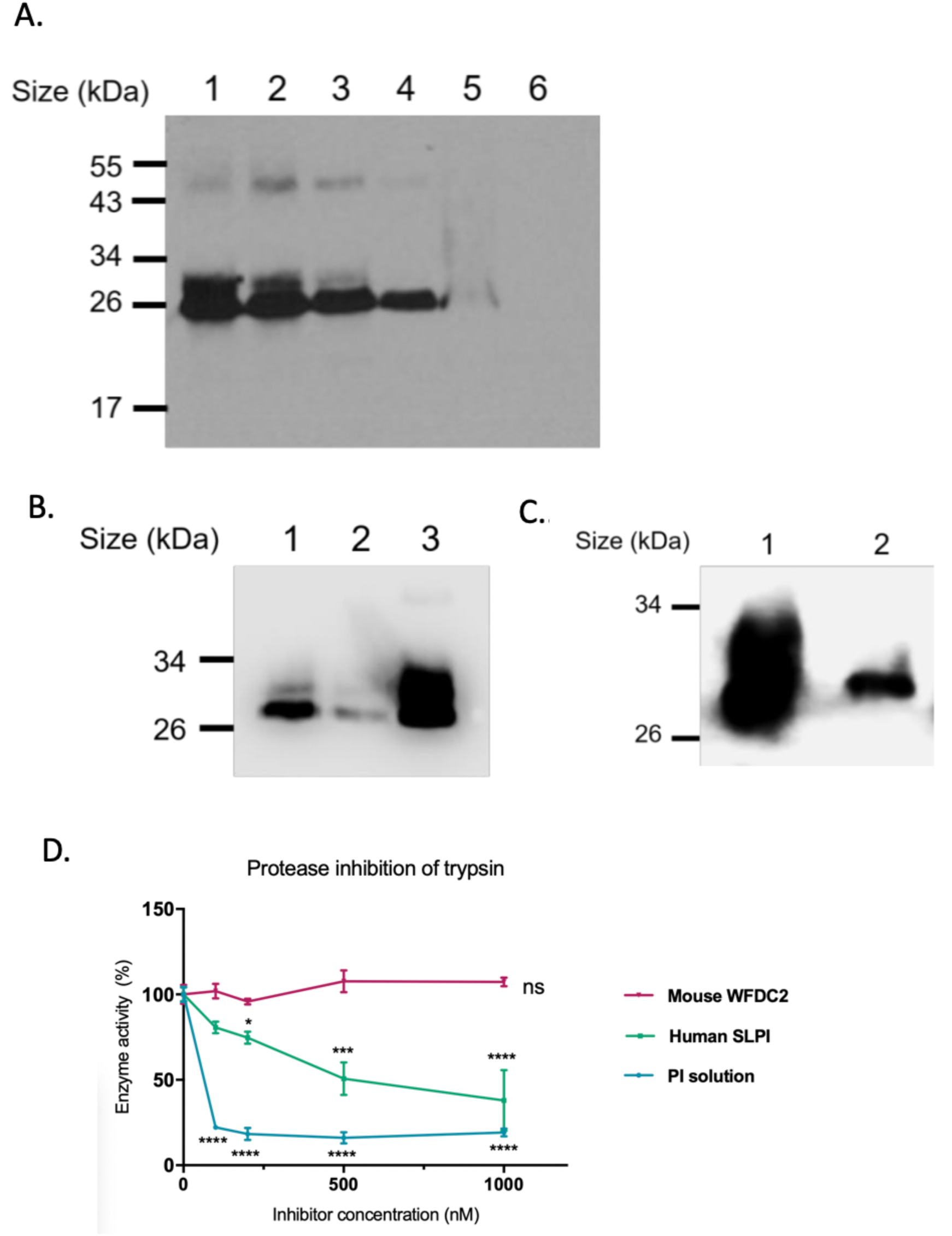
Functional studies of rWFDC2. **A.** HEK293 cells were transfected with pVR1255 WFDC2 serum-free conditioned media was collected 3 to 6 days post-transfection and analysed by Western blot using a polyclonal antibody against mouse WFDC2. Lane numbers: 1 – media collection day 3, 2 – media collection day 4, 3 – media collection 3 day 5, 4 – media collection day 6, 5 – transfected HEK293 cell lysate, 6 – conditioned media from cells transfected with empty pVR1255 vector. **B.** rWFDC2 was purified from conditioned media using a FLAG affinity resin and the eluted product analysed by Western blotting using anti-FLAG monoclonal antibody. Equal volumes of sample were loaded per lane. Lane numbers represent the following: 1 –conditioned media before purification, 2 –conditioned media after purification, 3 – purified protein. **C.** WFDC2 digested with sialidase was compared to untreated WFDC2 by Western blotting using anti-FLAG monoclonal antibody. Lane numbers represent the following: 1 – untreated WFDC2, 2 – Wfdc2 digested with sialidase enzyme (representative of n=3). **D**. The anti-trypsin activity of WFDC2, rSLPI and PI was determined as outlines in the Materials and Methods section. Results are expressed as a percentage reduction in enzyme activity with reactions without inhibitor representing 100% enzyme activity. Data represents the mean ± SD (n=3). Significance value is calculated as activity compared to enzyme without inhibitor; * P ≤ 0.01; *** ≤ 0.001; ****≤ 0.0001.

We used a chromogenic substrate assay to establish whether WFDC2 inhibited trypsin activity. Increasing amounts of WFDC2 protein were used (0-1000 nM) in the assay alongside recombinant human SLPI and a broad-spectrum protease inhibitor (PI) solution. The results show that murine WFDC2 had no significant effect on trypsin activity at concentrations as high as 1000 nM (Figure 6D) whereas the positive control SLPI caused up to a 50% decrease in enzyme activity and the broad-spectrum PI solution induced an 80% reduction in trypsin activity even at the lowest concentration tested (100 nM). Similar experiments undertaken with elastase also failed to show any inhibitory results (results not shown) and the protein also was incapable of inhibiting MMP2 and MMP9 (results not shown) in zymography assays.

The effect of WFDC2 on bacterial viability were assessed against *Streptococcus mutans, Streptococcus gordonii, Streptococcus pneumoniae* and *Escherichia coli*. In all assays, FLAG peptide elution solution was used as a control. The results show that the addition of 1uM mWFDC2 to bacterial culture had no effect on bacterial growth when compared to untreated bacteria, or bacteria incubated with FLAG peptide elution buffer or PBS (Figure 7A-D). Conversely, bacterial growth was significantly inhibited when cells were exposed to ampicillin (60 μg/ml). No change in absorbance was visible in wells loaded with sterile broth, showing that wells were not contaminated during incubation in the plate reader overnight. Increasing the amount of WFDC2 to 3um also had no effect (date not shown)

**Figure 7:**
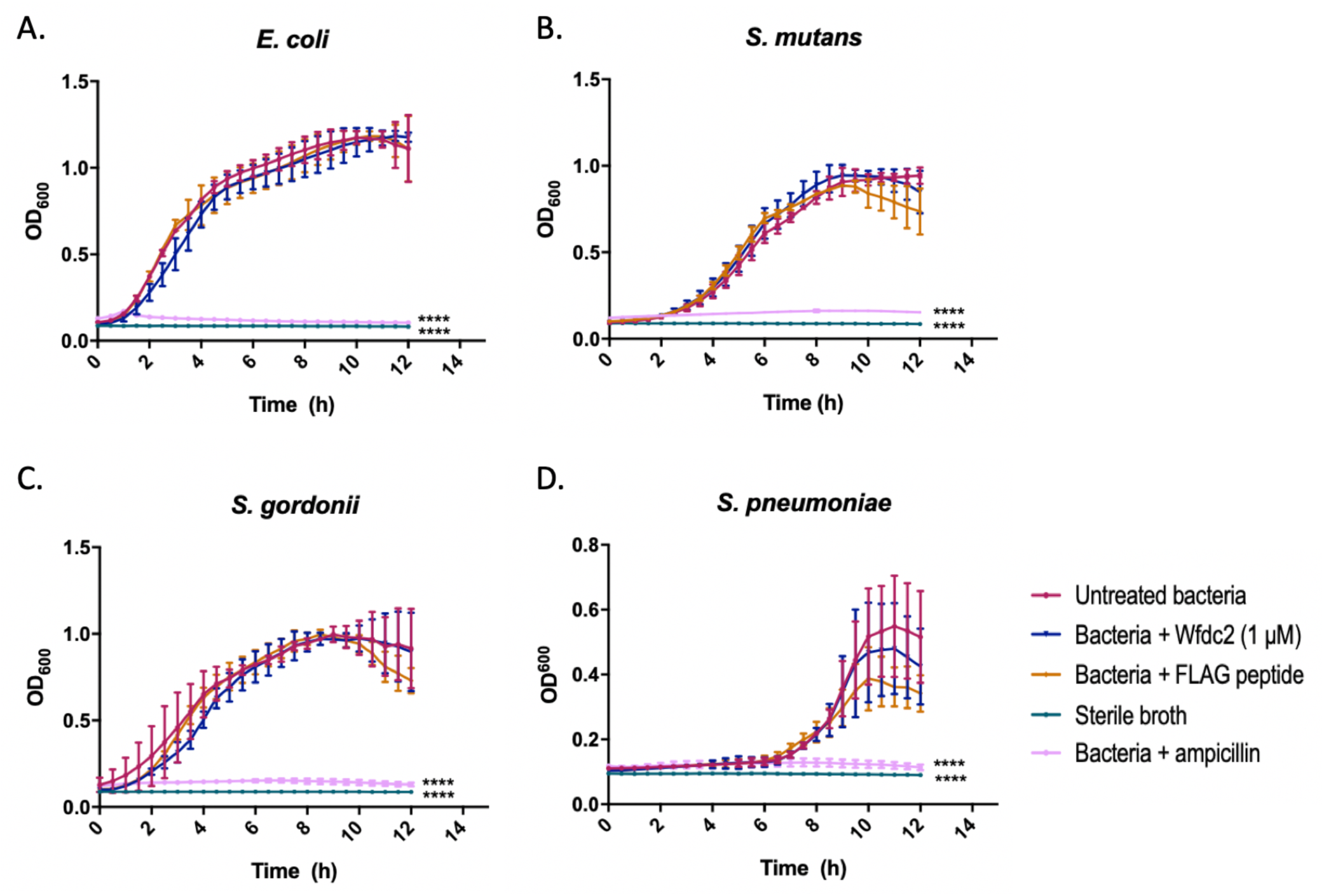
The effect of rWFDC2 on bacterial viability. Bacterial viability was measured using a growth curve technique as outlined in the Material and Methods section. Suspensions of (**A**) *E. coli*, (**B**) *S. mutans*, (**C**) *S. gordonii*, and (**D**) *S. pneumonia*, were incubated with rWFDC2 protein overnight. Changes in suspension optical density at OD^600^ were measured every 30 minutes for 12 hours using a plate reader to produce a growth curve. Data represents the mean ± SD (n=3); **** P ≤ 0.0001

## Discussion

WFDC2 was originally described as a small, secreted, epididymis-specific protein [12]. However, subsequent research has shown that this protein is also expressed in a range of tissues, including multiple tissues in the oral cavity as well as the respiratory and genital tracts [12, 32]. Previous studies of WFDC2 have focussed mainly on its use as a cancer biomarker due to an initial observation that is consistently overexpressed in ovarian cancer. More recently overexpression of WFDC2 has also been described in other tumours, including lung adenocarcinomas [17]. However, much of the biology of WFDC2 remains to be elucidated and its exact function remains unknown. Due to its similarity with other WFDC2 family members, it has been postulated that it may possess anti-protease and/or antimicrobial properties, as well as being involved in the regulation of inflammation and the immune response [11–13, 33].

The murine *wfdc2* gene contains an additional exon when compared to the majority of mammals. This exon, exon 3, is found between those encoding for the two WFDC domains and has the effect of separating the two domains. This unstructured linker is 53 amino acids in length. Our comparative analysis showed that this additional linker is only found in species from the *Cricetidae* and *Muridae* families of rodents and must have arisen since the divergence from the *Spalacidae* family (that includes the blind mole rat, *Nannospalax galili*). The linker region varies in length identified and is also very divergent. The sequence has no predicted structure and it seems likely that it serves to separate the two WFDC structural domains. The functional consequence of this additional sequence remains unclear. The consequence of the alternative splicing of murine WFDC2 also awaits clarification. We previously reported that human WFDC2 undergoes alternative splicing to potentially yield a number of distinct protein isoforms [12,34] and it has been suggested that expression of different amounts of these may have some diagnostic or prognostic value in cancers [34]. In both species, alternative splicing can potentially generate protein isoforms containing just a single WFDC domain. The mouse gene can also potentially yield an isoform that lacks the linker region (encoded by exon 3) and is, therefore, similar to the proteins found in the majority of other species. Specific analysis of the distinct isoforms will be needed to establish if any exhibit differential expression and analysis of the distinct protein products will be required to confirm their distinct functions.

Our results clearly demonstrate that *wfdc2* is readily detectable in a range of tissues from the mouse head and oral cavity as well as in the respiratory tract. The highest levels of expression appear to be in the upper respiratory tract and regions of the nasal passages. In general, the distribution of the protein within the mouse respiratory tract, oral cavity and nasal passage tissue is similar to that reported in man [12, 13, 16, 32]. In the major salivary glands, with the exception of the submandibular gland, the protein is expressed by ductular cells rather than by the acinar cells. In the submandibular gland there is strong expression in the mucous tubules. The localisation of the protein within the major glands is consistent with reports that the protein is secreted into mouse saliva [35]. The protein is also found in a small number of minor mucosal glands in the head and neck, suggesting that it is constitutively secreted into the oral cavity. The distribution within the epithelial cells of the respiratory tract and head regions is of some interest. Our IHC data does not suggest that the protein is restricted to a specific cell type but rather that it is found in a number of different cells. In some cases, for example within the nasal septum and conjunctiva, the protein is found in a population of goblet cells whereas in the lung the protein stains strongly in the bronchiolar epithelial cells that appear to represent club (Clara) cells. In other locations, for example in the epithelium of the epiglottis, staining is seen in the squamous epithelium. This lack of cell type specific localisation is consistent with recent data generated by scRNA seq which shows gene expression in multiple epithelial cell types throughput the lung (Figure S2). These observations suggest that the molecular regulation of *wfdc2* in these regions is governed by different mechanisms to those reported for cell type specific genes.

Our data shows that mouse WFDC2 is secreted into mucosal fluids in the lung and mouth. These observations are consistent with those seen in man, where the protein has been reported in nasal secretions, bronchioalveolar lavage fluid, saliva, tears and endocervical mucus [36–40]. It has been identified in murine nasal secretions [41] and saliva [35]. The protein appears as multiple bands in mouse BAL fluid samples and contains multiple potential O-glycosylation sites. Unlike the human protein [16] mouse WFDC2 is not N-glycosylated.

Given that murine WFDC2 has a similar distribution to that seen in man, what might the functional implications be? Because of the sequence similarity between WFDC2 and other WFDC-domain containing proteins, principally SLPI, it has been thought that the protein would function as an antiproteinase [11,13]. However, there is only limited supportive data in the literature [21, 33] suggesting perhaps the protein functions as an immune regulator and is part of the innate immune defence shield protecting mucosal surfaces, a role similar to that identified for SLPI [10–12]. To date, WFDC2 has been reported to have limited antimicrobial activity [21,22, 33] but has been shown to bind to bacteria [42,43] and perhaps serves to ensnare pathogens as part of the mucosal innate immune response. In our study using recombinant mouse WFDC2, we were unable to identify any proteinase inhibitory activity nor were we able to show robust antimicrobial activity against a number of bacteria. Other studies, largely from the cancer field, have shown that WFDC2 has the capacity to promote cell growth [43–48].

Recently, independent *Wfdc2* deficient mouse lines have been generated and shown to have an embryonic lethal phenotype [23, 24]. *Wfdc2*^*-/-*^ mice die at, or soon after, birth from respiratory failure [23, 24]. *Wfdc2*^*-/-*^ newborn mice exhibit atelectasis and unsurprisingly exhibit a significant enhanced inflammatory profile, that is lacking from lungs preterm [23]. This is associated with phenotypic alterations of the airway epithelium in the developing *Wfdc2*^*-/-*^ lung [23] and μCT analysis shows a reduction in luminal diameter of the major airways prior to birth (unpublished). These observations are consistent with a delay in lung development. Robust *Wfdc2* expression is seen throughout lung development [23, 49] and this data suggest that the protein functions somehow to promote lung development. Roles for WFDC2 outside of the wider respiratory tract remain unknown.

---

**Supplementary Figure. 1.**
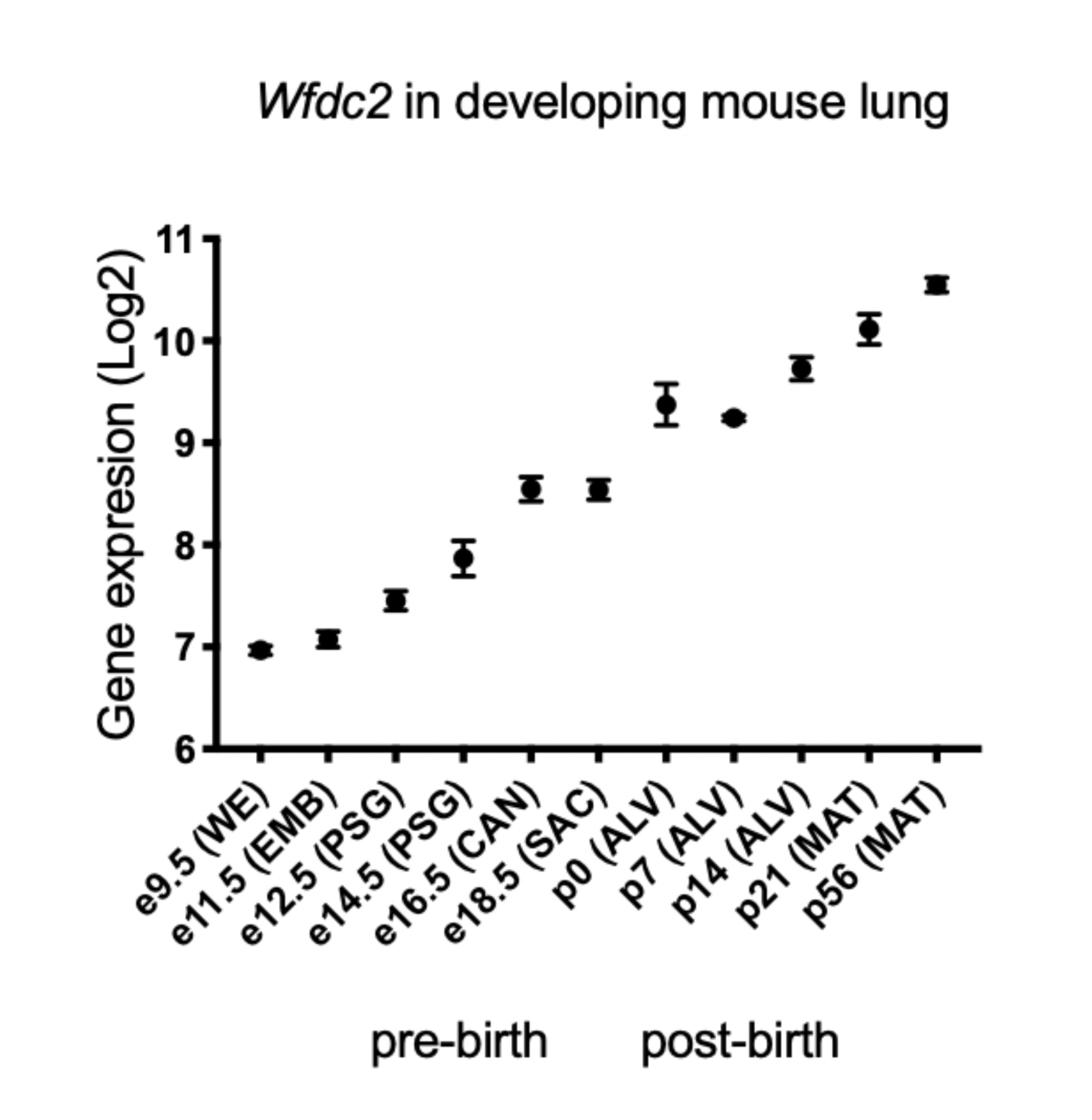
*Wfdc2* expression steadily increases during mouse lung development. Expression data for *Wfdc2* in C57/BL6 mouse lungs during development [50] was extracted from array data in LungGENs [51, 52]and gene expression plotted as Log2 expression using Prism. Data is presented as mean + SEM (n=4-6).

**Supplementary Figure. 2.**
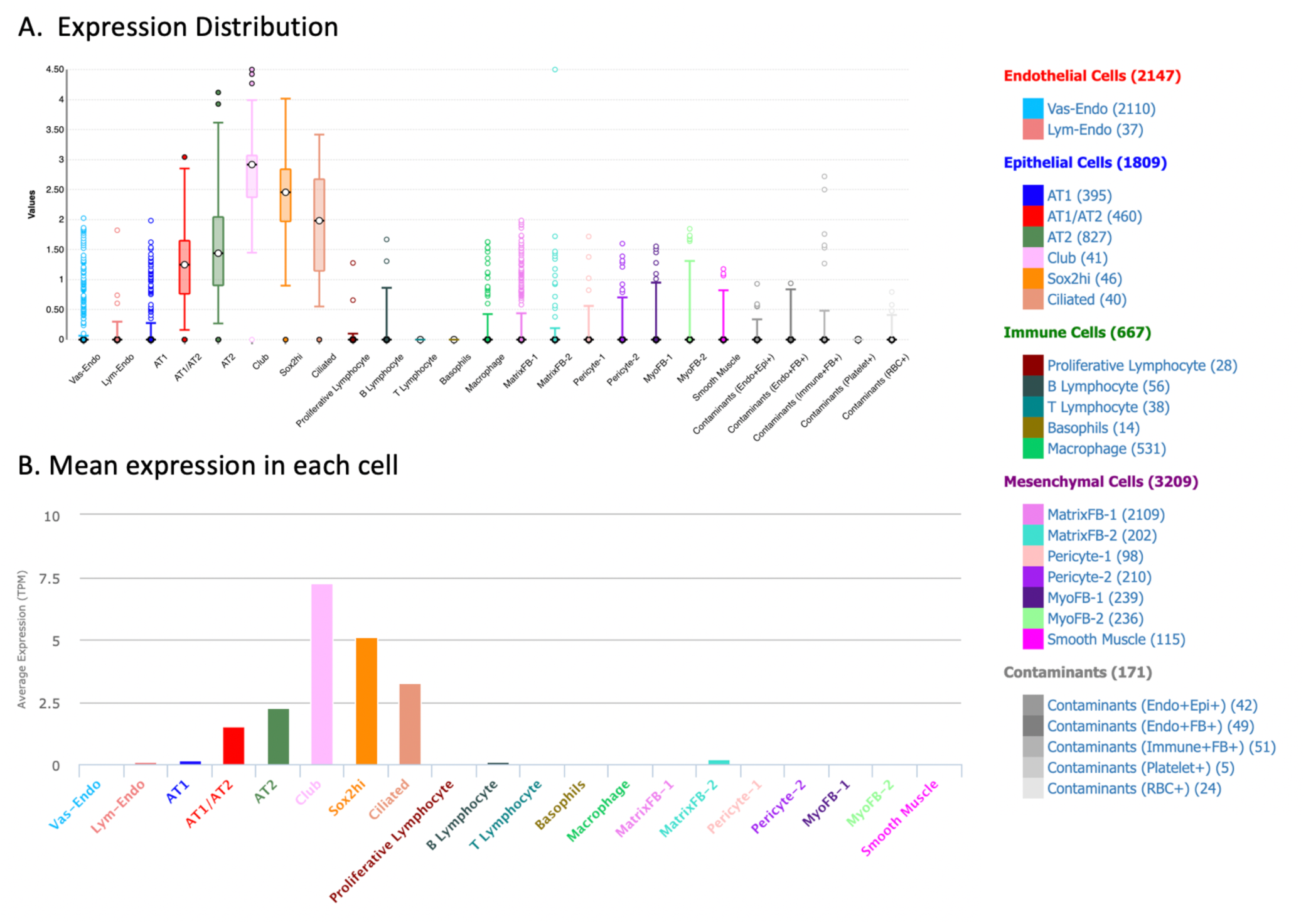
Wfdc2 is expressed in multiple epithelial cell types within the mouse lung. Cell type specific expression of *Wfdc2* in post-natal day 1 mouse lung was extracted from LungGENS [51,52]. Data can be found on https://research.cchmc.org/pbge/lunggens/genequery_dp.html?spe=MO&tps=pnd1&geneid=Wfdc2) **A.** shows expression distribution. **B.** shows mean expression in each cell. Data was from over 8000 cells from two individual mice.

## Acknowledgements

We acknowledge the help of Helen Marriott in providing the oral wash and bronchoalveolar lavage samples, Apoorva Mulay for providing the dissected nasal turbinate tissue samples and culturing the mNEC and mMEEC cultures and Priyanka Anjuan for providing the mTEC cultures. We thank Dr Paul Heath for running the DNA arrays.

